# Somatic nuclear blebbing may indicate proliferating germ cells but does not indicate organismal aging in *Caenorhabditis elegans*

**DOI:** 10.1101/2022.09.17.508355

**Authors:** Qiang Fan, Xue-Mei Li, Chao Zhai, Shang-Tong Li, Meng-Qiu Dong

## Abstract

Abnormal nuclear morphology is suggested to be a hallmark of aging. One type of such abnormalities is nuclear blebbing, but little is known about whether and how nuclear blebbing participates in animal aging. What regulates nuclear blebbing is also unknown. In this study, we show that the frequency of nuclear blebbing in the hypodermis increases during aging in wild-type *C. elegans*. These nuclear blebs are enveloped by the nuclear lamina, the inner and the outer nuclear membrane, and 42% of them contain chromatin. Detachment of a bleb from the nucleus is rare but does happen, thereby generating cytoplasmic chromatin. Cytoplasmic chromatin-containing lysosomes juxtaposing the nucleus are detected in old worms. Therefore, nuclear blebbing contributes to the age-associated chromatin loss. However, the frequency of nuclear blebbing does not correlate with the rate of aging in *C. elegans*. Old age does not necessarily induce nuclear blebbing, neither does starvation, heat stress, or oxidative stress. Intriguingly, we find that proliferation of germ cells promotes nuclear blebbing.

**Highlights:**

i. Nuclear blebs accumulate in the hypodermis during *C. elegans* aging
ii. Nuclear blebbing contributes to chromatin loss
iii. The frequency of nuclear blebbing does not correlate with the rate of aging
iv. Proliferating germ cells promote nuclear blebbing in the hypodermis

## 1. Introduction

In eukaryotic cells, the nucleus is enclosed by the nuclear envelope, which consists of two lipid bilayer membranes, the outer nuclear membranes (ONM) and the inner nuclear membranes (INM). The two membranes are fastened up by the <u>Li</u>nker of <u>N</u>ucleoskeleton and <u>C</u>ytoskeleton (LINC) complexes. LINC complexes are composed of KASH proteins anchored to ONM and SUN proteins anchored to INM. INM is lined with the nuclear lamina, a meshwork of mostly lamins, which bind to each other and also to INM proteins such as emerin (Cohen-Fix and Askjaer, 2017). Scattered over the nuclear envelope are nuclear pores. Built up from hundreds of individual proteins, the megadalton nuclear pore complexes (NPC) span across ONM and INM to allow material exchange between the nucleoplasm and the cytoplasm. Together, these proteins maintain the normal structure and function of the nucleus.

Normally, nuclei are oval-shaped, but abnormalities in nuclear morphology arise during aging (Scaffidi and Misteli, 2006; Schreiber and Kennedy, 2013; Zink et al., 2004) and abnormal nuclear morphology is suggested to be a hallmark of aging (Pathak et al., 2021). Convolution of the nuclear membrane or nuclear lamina, detected in most cases using a fluorescently labeled lamin protein, is a visually striking, gross abnormality of the nuclear shape and is referred to here as nuclear membrane convolution for simplicity. Nuclear membrane convolution is a characteristic of fibroblasts isolated from normally aged humans (Scaffidi and Misteli, 2006). In aged *C. elegans*, the nuclear membrane also becomes convoluted (Haithcock et al., 2005). Furthermore, age-associated nuclear membrane convolution is slowed down in the *daf-2(e1370)* mutant (Haithcock et al., 2005; Zhao et al., 2017) and the *eat-2(ad1116)* mutant (Charar et al., 2021), both are long-lived compared to the wild type worm. However, pharmacological inhibition of farnesylation of lamin proteins, which ameliorates age-associated nuclear membrane convolution, fails to extend *C. elegans* lifespan (Bar and Gruenbaum, 2010; Bar et al., 2009). This suggests that nuclear membrane convolution can be uncoupled from aging. However, other types of nuclear abnormalities such as nuclear blebbing have not been examined from this perspective.

Relative to nuclear membrane convolution, which is a global deformation of the nucleus, nuclear blebbing is a localized deformation, which involves the formation of relatively small protrusions from the surface of a nucleus. Nuclear blebs can be seen on otherwise smooth, oval-shaped nuclei of young adult animals. It has been reported that nuclear blebs in cultured senescent cells can mediate the nucleus-to-cytoplasm transport of chromatin and lamin B1 (Dou et al., 2017; Dou et al., 2015; Ivanov et al., 2013). The abnormal presence of chromatin in the cytoplasm activates the cGAS-STING (cyclic GMP–AMP synthase linked to stimulator of interferon genes) pathway, which senses cytosolic DNA and in turn, promotes secretion of pro-inflammatory cytokines, a key feature of senescent cells (Dou et al., 2017). Additional studies show that ectopic lamin B1 in the cytoplasm is targeted to the lysosome for degradation, and the loss of lamin B1 promotes cellular senescence (Dou et al., 2015; Ivanov et al., 2013). Thus, nuclear blebbing may mediate cellular senescence.

Nuclear blebs have been observed during aging in *C. elegans*, but remain poorly characterized (Haithcock et al., 2005). Using *C. elegans* as an aging model, we try to find out in this study: 1) whether the nuclear membrane surrounding a nuclear bleb is of an intact structure with normal ONM, INM, nuclear lamina and nuclear pores; 2) whether chromatin is present inside a nuclear bleb and whether blebbing leads to degradation of chromatin in the cytoplasm as has been reported in cultured cells; 3) how nuclear blebbing changes with age; and 4) whether nuclear blebbing is coupled with organismal aging. Below we report our findings, which answer these questions, and an additional intriguing discovery that nuclear blebbing in the hypodermis responds to proliferating germ cells in the gonad.

## 2. Methods

### 2.1 *C. elegans* strains

The following strains were used in this work:

UD484 (yc32[gfp::lmn-1] I) (Bone et al., 2016)
MQD2907 (yc32[gfp::lmn-1] I; bqSi226[emr-1p::emr-1::mCherry + unc-119(+)] IV)
MQD2908 (bqSi226[emr-1p::emr-1::mCherry + unc-119(+)] IV; hqIs466[npp-6::gfp])
MQD2615 (yc32[gfp::lmn-1] I; thu7[his-72::mcherry])
MQD1844 (p720–4[lmn-1p::emr-1::gfp::unc-54 3’UTR + unc-119(+)]; thu7[his-72::mcherry])
MQD2029 (thu7[his-72::mcherry]; qxIs430[scav-3p::scav-3::gfp])
MQD1807 (thu7[his-72::mcherry]; qxIs520[Pvha-6::laat-1::gfp])
MQD2808 (yc32[gfp::lmn-1] I; daf-2(e1370) III)
MQD2917 (glp-1(e2141) III; yc32[gfp::lmn-1] I)
MQD2918 (fer-15(b26ts) II; yc32[gfp::lmn-1] I)
LW699 (p720–4[lmn-1p::emr-1::gfp::unc-54 3’UTR + unc-119(+)]) [11]
MQD2423 (bqSi235 [emr-1p::emr-1::GFP + unc-119(+)] II)
MQD2425 (bqSi226 [emr-1p::emr-1::mCherry + unc-119(+)] IV)
MQD2684 (hqKi450[emr-1::gfp] I)

### 2.2 plasmid

Plasmids used to construct EMR-1::GFP knock in strain:

EGKI-HR (carrying repair template for homology directed repair, modified from pPD95_77)
EGKI-sg1 and EGKI-sg14 (carrying sgRNA, modified from pDD162)

Plasmids used to construct GFP::GFP::KASH knock in strain, GFP::GFP::KASH was inserted into the single copy insertion site on chr II:

GGKKI-HR (carrying repair template for homology directed repair, modified from pPD95_77)
GGKKI-sg1 and GGKKI-sg2 (carrying sgRNA, modified from pDD162)

### 2.3 Imaging and measurement of nuclear blebbing frequency

Worms were cultured under standard conditions, i.e., 20°C on NGM plates seeded with live OP50. Images were acquired with a spinning-disk confocal microscope (UltraVIEW VOX; PerkinElmer) equipped with a 63 ×, 1.4 numerical aperture oil-immersion objective. The exposure time and laser power were varied to balance the fluorescence intensity among samples. Nuclear blebs were identified manually.

### 2.4 Stress condition

For heat stress, on AD2, worms were exposed to 37°C for 2 h and imaged immediately; for starvation stress, on AD2, worms were transferred to NGM plates containing no OP50 and imaged 10 hours after starvation initiation; for paraquat (PQ) stress, worms were hatched and cultured on NGM plates containing live OP50 and 0.1 mM or 2 mM PQ and imaged on AD3.

### 2.5 Scanning electronic microscopy analysis

EM samples and Scanning EM imaging were conducted according to the methods developed by Li et al (Li et al., 2017). Worms at adult day 2, day 9, and day 18 were collected and were fixed via high-pressure freezing (Wohlwend HPF Compact-01). Fixed samples were put into 1% OSO4 acetone solution for dehydration (Leica AFS2), and the temperatures control program was set as −90□ for 72 hrs.; increase 2□ per hour until reach −60□; −60□ for 10 hrs.; increase 2□ per hour until reach −30□; −30□ for 10 hrs.; increase 2□ per hour until reach 4□. Staining samples with UAc saturated acetone solution for 3.5 hrs. at room temperature. Use SPI-PON812 resin (SPI-CHEM) for filtration, then samples were embedded and polymerized at 60□ for 48 hrs. 70 nm sections sectioned by ultramicrotome (UC6, Leica, Germany) were collected on PE tap (Shanghai Jinghou Electronics Technology Co., Ltd. China) and stuck on a silicon wafer (Suzhou crystal silicon electronic & technology Co., Ltd. China) via electroconductive adhesive tape (NISSHIN EM Co. Ltd, Japan). Sections were examined via SEM (FEI Helios NanoLab 600i) equipped with CBS detector with 2.0 kV in accelerating voltage, 0.69 nA in current, 10 μs in dwell time, and visualized via the software, xT microscope control (FEI, version 5.2.2.2898) and PinPoint (developed by Li et al. (Li et al., 2017)).

### 2.6 Real-time PCR

At each time point, 50 worms were collected into 50 μl of lysis buffer and frozen immediately in liquid nitrogen. Lysis was carried out by incubation at 60°C for 1 h and proteinase K was inactivated by incubating at 95°C for 10 min. RT-PCR was carried out on an ABI 7500 Fast Real-Time PCR System using a TAKARA real-time PCR kit (SYBR Premix Ex TaqTM II). The primers used were as follows:

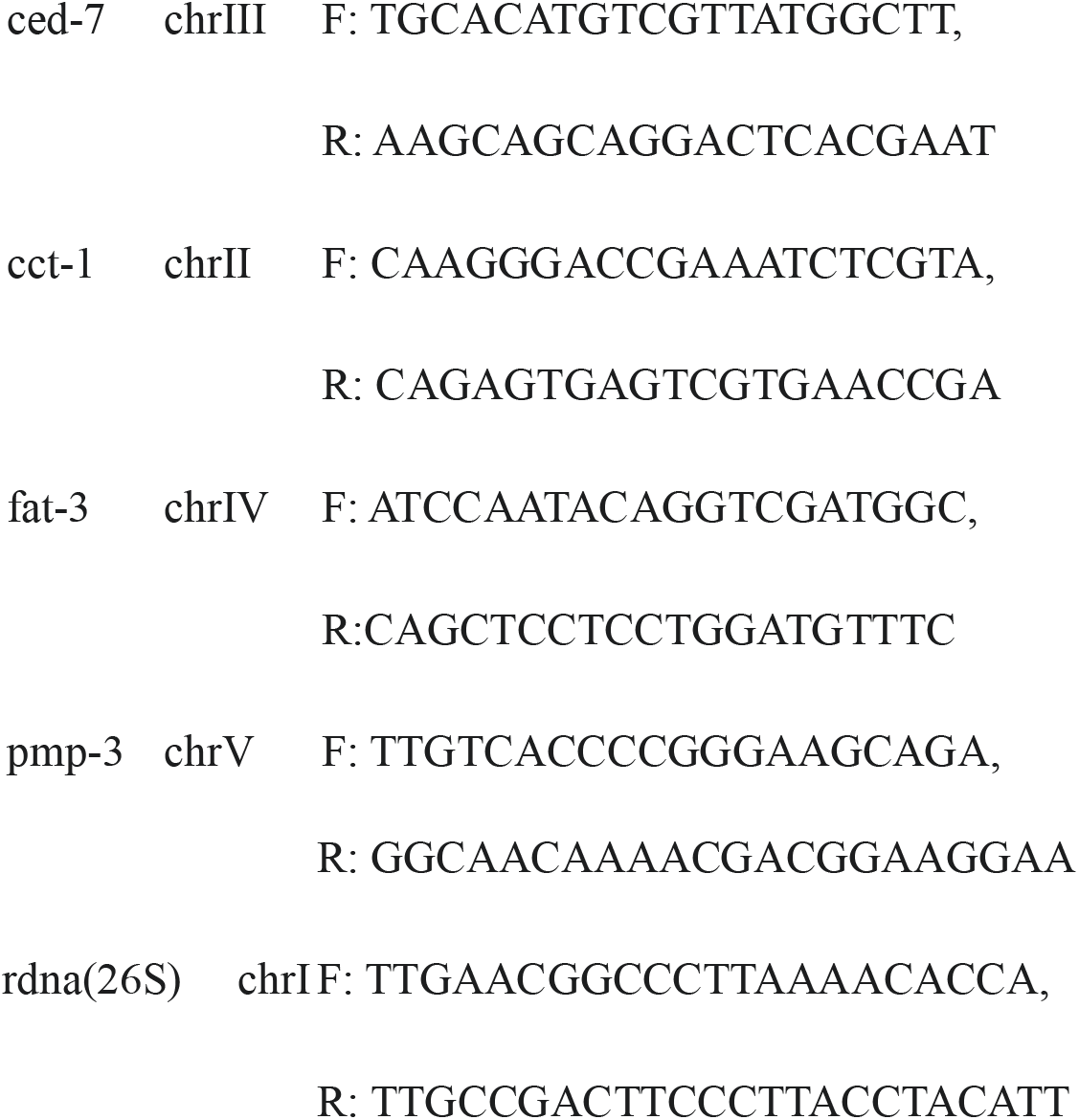

### 2.7 Quantitative western blot analysis

For measuring H3 protein levels, *glp-1(e2141)* mutants were hatched and cultured at 25 °C. At each timepoint, 100 worms were collected into 20 μL M9 buffer, flash-frozen and stored at −80 °C. The worms were thawed, mixed with 20 μL 2x SDS loading buffer, boiled for 15 min at 100 °C before loading. Blots were incubated with anti-H3(Abcam, 1791), 1:5000. Quantification was carried out using ImageJ.

### 2.8 γirradiation

For irradiation, on AD2, worms were exposed to 80 Gy irradiation using a Gammacell 1000 blood irradiator and imaged 10 hours later.

### 2.9 FUDR treatment

For FUDR treatment, 6-cm NGM plates containing 100 μM FUDR were seeded with 200 μL concentrated OP50. Worms were transferred to FUDR plates since mid L4 stage.

### 2.10 ThT treatment

For ThT treatment, 6-cm NGM plates containing 100 μM ThT were seeded with 200 μL concentrated OP50. Worms were hatched and cultured on ThT plates.

## 3. Results

### 3.1 Nuclear blebbing frequency increases during aging in wild-type *C. elegans*

We started out by characterizing nuclear blebs in aged *C. elegans*. We chose to focus on hypodermal nuclei, specifically, the nuclei of the hyp7 cell, which wraps around most of the worm body. Large and thin, hyp7 has a total of 139 nuclei, a result of extensive cell-cell fusion during embryonic and larval development (Shemer and Podbilewicz, 2000). The hyp7 nuclei are relatively large, uniform in appearance, and close to the body surface, thus yield high image quality. LMN-1 is the sole lamin protein in *C. elegans*. By imaging a knock-in worm strain expressing GFP::LMN-1 from the endogenous *lmn-1* locus, we observed that hyp7 nuclei from young animals had few blebs whereas those from aged worms had many (Fig 1A). The frequency of nuclear blebbing increased with age, from close to zero on adult day 1 (AD1) to over 30 blebs per 100 hyp7 nuclei on AD14 (Fig 1B). Neither heat stress, starvation, nor paraquat treatment, which produces reactive oxygen species (ROS) in the cell (Shen et al., 2014), increased the frequency of nuclear blebbing (Fig 1C), suggesting that nuclear blebbing is a rather specific phenomenon associated with old age, not something induced by stress in general.

**Fig 1.**
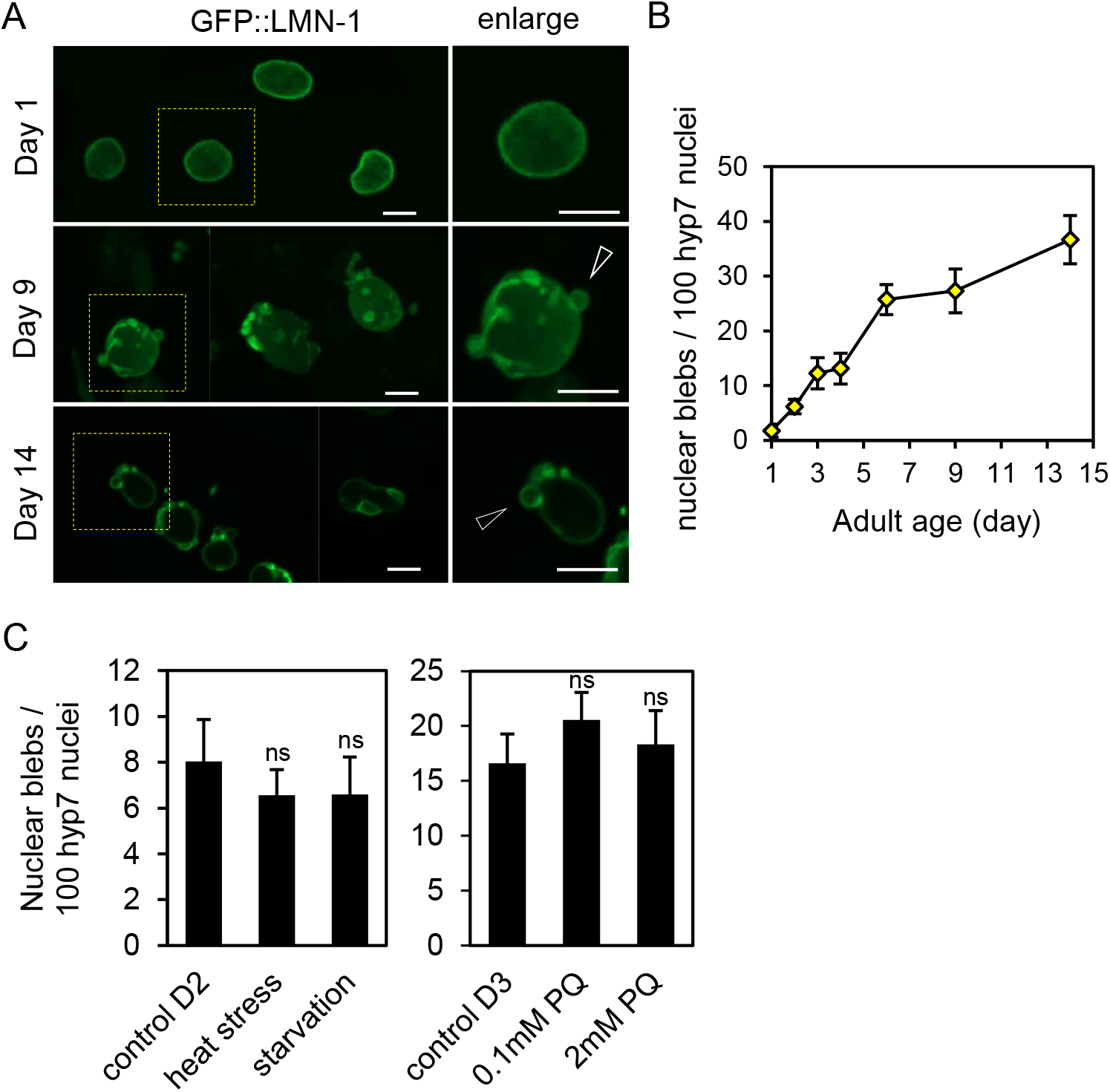
Nuclear blebs accumulate during aging in *C. elegans*. (A) Nuclei of hyp 7 on adult day 1, 9, and 14. The boxed regions are magnified and shown on the right, with arrowheads pointing to nuclear blebs. (B) Age-dependent increase of nuclear blebs. (C) The indicated stress conditions did not induce nuclear blebbing. Adult day 2 (D2) worms were subjected to heat stress and starvation, and adult day 3 (D3) worms were treated with paraquat (PQ). ns, not significant, as determined by t-test. Error bars represent standard error. Scale bars represent 5 μm. The hyp 7 nuclei of the *gfp::lmn-1* knock-in strain UD484 were imaged and analyzed in (A-C).

### 3.2 Characteristics of nuclear blebs

Before examining the relationship between nuclear blebbing and *C. elegans* aging, we asked whether nuclear blebs are enveloped by normal nuclear membrane. We used EMR-1::mCherry to mark INM and examined it with respect to GFP::LMN-1, and found that they colocalized in all hyp7 nuclear blebs examined (Fig 2A). Nuclear blebs were also positive for GFP::GFP::KASH, a marker of ONM with a tandem GFP tag (Fig 2B). GFP::LMN-1, EMR-1::mCherry, and GFP::GFP::KASH signals were all enriched on the periphery of nuclear blebs, just like the mother nuclei from which they protrude (Fig 2A-B). Electron microscopy (EM) analysis confirmed the presence of ONM and INM in nuclear blebs (Fig 2C). We then asked whether nuclear blebs also possess nuclear pores as the mother nuclei do. NPP-6, the *C. elegans* ortholog of human nucleoporin 160, is a peripheral subunit of NPC of multiple copies (16 copies per NPC) (Tie et al., 2016). We tagged NPP-6 with GFP to label nuclear pores and found that 78% of the nuclear blebs (*n* = 37) marked by EMR-1::mCherry contained no NPP-6::GFP, suggesting that nuclear blebs are deficient in NPCs or nuclear pores (Fig 2D).

**Fig 2.**
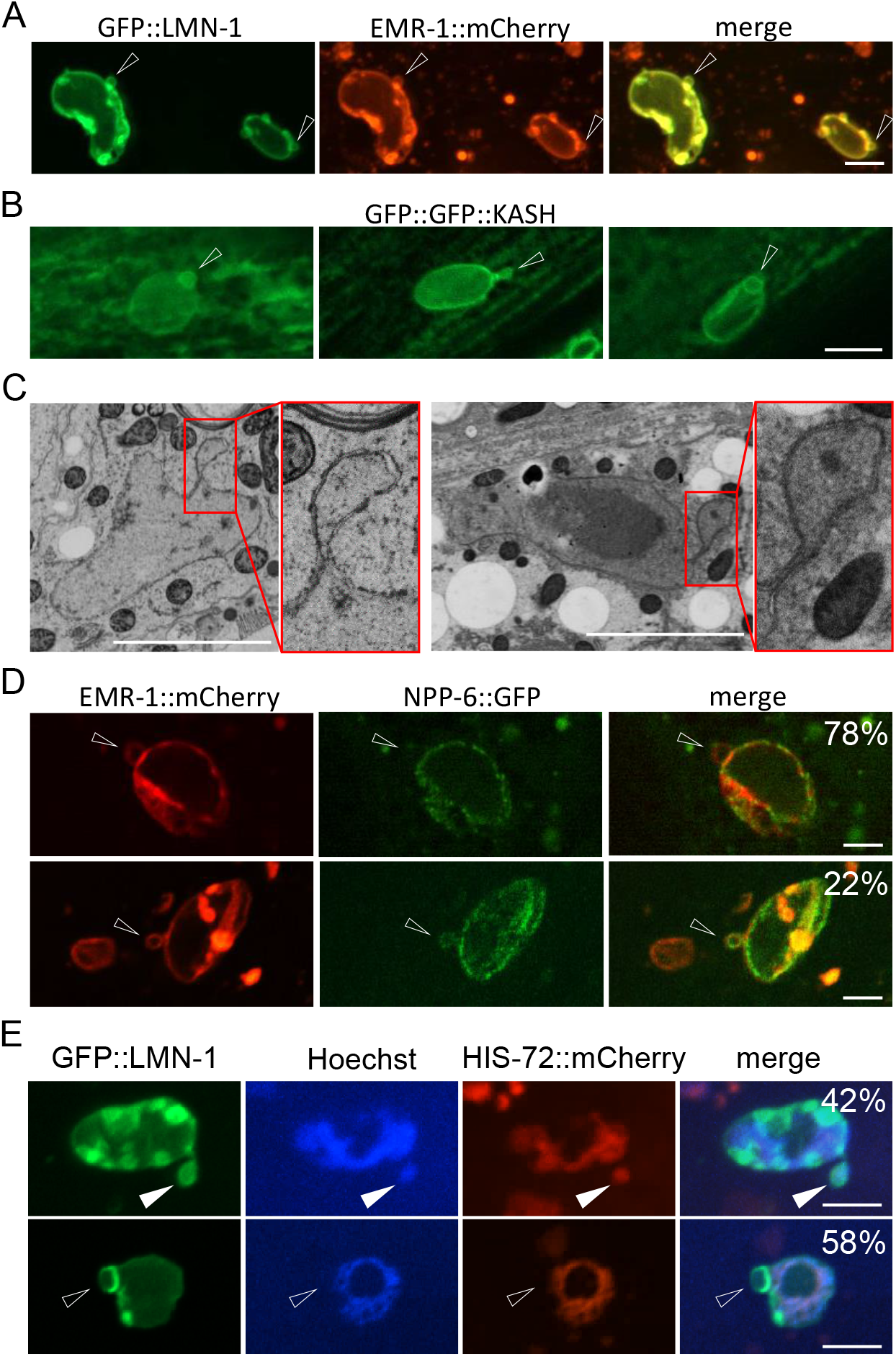
Characteristics of nuclear blebs in aged *C. elegans*. (A) Double labeling of nuclear blebs by a nuclear lamina marker GFP::LMN-1 and an INM marker EMR-1::mCherry. (B) Labeling of nuclear blebs by an ONM marker containing a tandem GFP tag fused with the KASH domain of UNC-83. (C) Two EM images of nuclear blebs. Selected regions (smaller boxes) are magnified and shown as insets (larger boxes) (D) 78% nuclear blebs marked by EMR-1::mCherry were not labeled by NPP-6::GFP (upper row), but 22% were (lower row). (E) On adult day 4, 42% of the nuclear blebs contained chromatin (upper, filled arrowhead) and 58% not (lower, empty arrowhead). Scale bars, 5 μm.

We further asked whether nuclear blebs contain chromatin. Using HIS-72::mCherry to indicate the presence of histones and the Hoechst dye to mark DNA, we found that 42% of the nuclear blebs contained both HIS-72::mCherry and DNA, whereas 58% had neither (*n* = 45) (Fig 2E). This intriguing result indicates that only less than 50% of nuclear blebs contain chromatin.

In summary, we find that *C. elegans* hyp7 nuclei gradually form and accumulate nuclear blebs as the animal grows old. These blebs have ONM, INM, and the nuclear lamina, but only 42% of them contain chromatin.

### 3.3 Nuclear blebbing contributes to chromatin loss in aged *C. elegans*

Next, we asked whether nuclear blebbing leads to chromatin loss. Using time-lapse imaging, we tracked more than 10 nuclear blebs for 8 h and found one that detached from the mother nucleus during the observation period (Fig 3A). Thus, although nuclear blebbing can give rise to cytoplasmic chromatin, such events are rare in *C. elegans*. To find out whether lysosomes degrade cytoplasmic chromatin in *C. elegans* as has been reported in cultured senescent cells (Ivanov et al., 2013), we examined the colocalization of cytoplasmic chromatin with two fluorescently labeled lysosomal membrane proteins SCAV-3 and LAAT-1. We indeed found lysosomes positive for both HIS-72::mCherry and SCAV-3::GFP or LAAT-1::GFP positioned right next to the hyp7 nuclei (Fig 3B and 3C). The size and the position of such histone containing lysosomes invite comparison with detached nuclear blebs.

**Fig 3.**
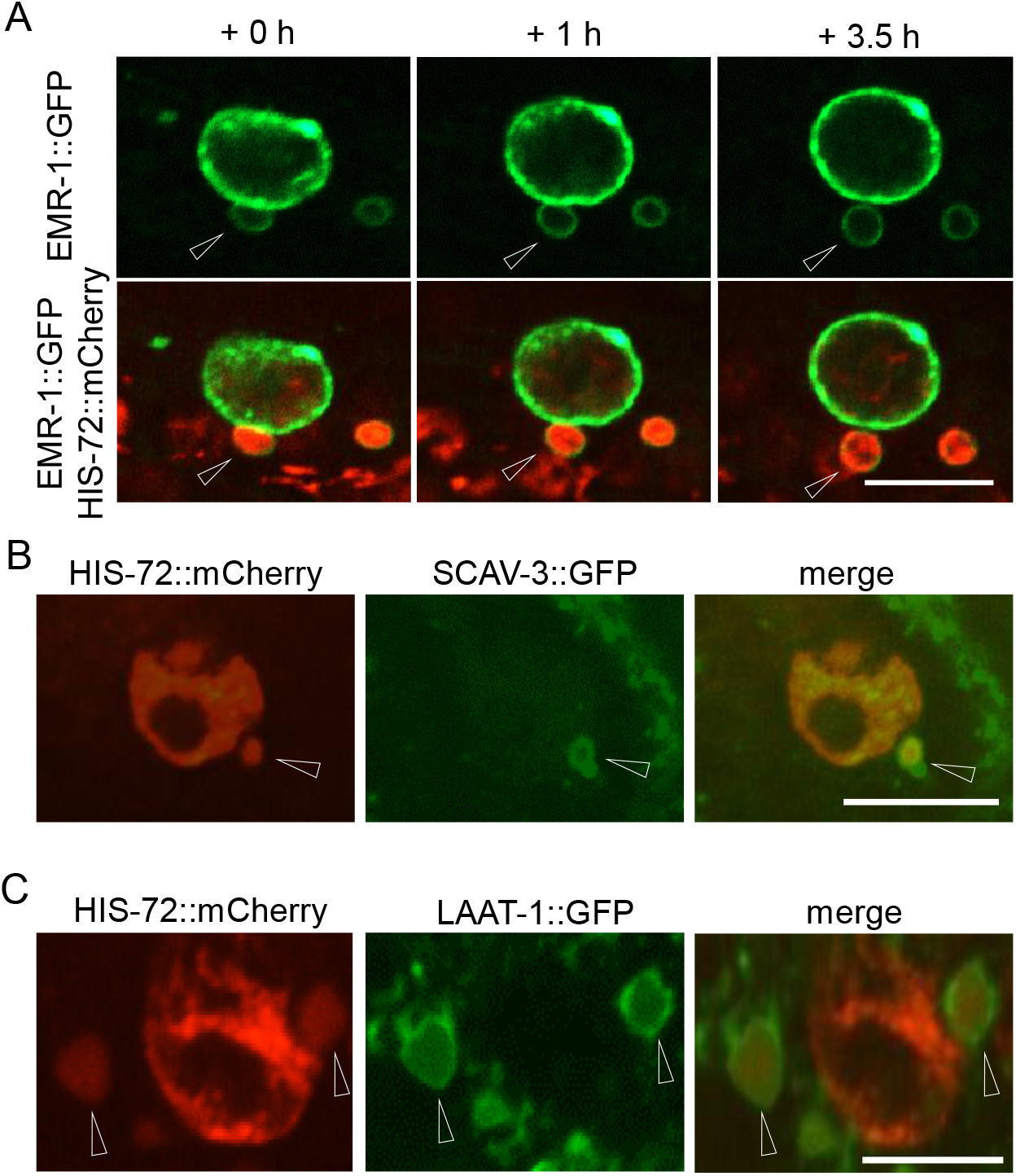
Nuclear blebbing contributes to chromatin loss. (A) Detachment of a nuclear bleb dually labeled by EMR-1::GFP and HIS-72::mCherry. (B, C) A subpopulation of cytoplasmic chromatin was found in lysosomes marked by either SCAV-3::GFP (B) or LAAT-1::GFP (C). Arrowheads indicate nuclear blebs. Scale bars, 5 μm.

Also consistent with the idea that nuclear blebbing can lead to chromatin degradation, it has been shown that the DNA copy numbers per worm decreased with the age of the animal (Golden et al., 2007). Here we repeated the experiment and verified the finding on each of the five representative genes (one encoding rRNA and four coding proteins, each on a different chromosome) (Fig S1A and (Golden et al., 2007)). The protein level of histone H3 also decreased gradually during aging (Fig S1B).

Next, we considered a plausible hypothesis that nuclear blebbing may mediate degradation of damaged chromatin. Previous studies have shown that nuclear blebs in cultured cells are enriched with γ-H2AX, a classic marker of damaged chromatin (Ivanov et al., 2013; Karoutas et al., 2019). As the *C. elegans* genome does not encode γ-H2AX, we did not examine directly whether damaged chromatin was enriched in nuclear blebs. Instead, we induced DNA damage by subjecting *C. elegans* to ionizing irradiation and expected the frequency of nuclear blebbing to spike after irradiation if nuclear blebbing were a significant mechanism for the animal to remove damaged chromatin. However, this was not the case (Fig S1C).

Hence, the above results demonstrate that in aged *C. elegans*, nuclear blebbing sporadically leads to the severance of chromatin from the nucleus, giving rise to cytoplasmic chromatin, which is then targeted by the lysosome for degradation. However, our data do not support the idea that nuclear blebbing mediates the nucleus-to-cytoplasm transport of damaged chromatin.

### 3.4 The frequency of nuclear blebbing does not correlate with the rate of aging in *C. elegans*

Next, we asked whether nuclear blebbing is modulated by, or even modulates, the rate of aging in *C. elegans*. The insulin/IGF-1-like signaling pathway regulates aging across animal species (Kenyon, 2010). The *daf-2(e1370)* mutant *C. elegans*, in which the insulin/IGF-1 receptor gene is compromised, lives twice as long as the wild type (Kenyon et al., 1993). However, in this long-lived *daf-2(e1370)* mutant, the frequency of nuclear blebbing is hardly different from that in wild-type worms for as long as two weeks into adulthood (Fig 4A). Temperature is a key environmental factor that controls lifespan of *C. elegans*. At elevated temperatures such as 25 °C, worms age faster (Hosono et al., 1982). However, a 5 °C increase or decrease from the standard culture temperature of 20 °C did not accelerate or decelerate nuclear blebbing (Fig 4B).

**Fig 4.**
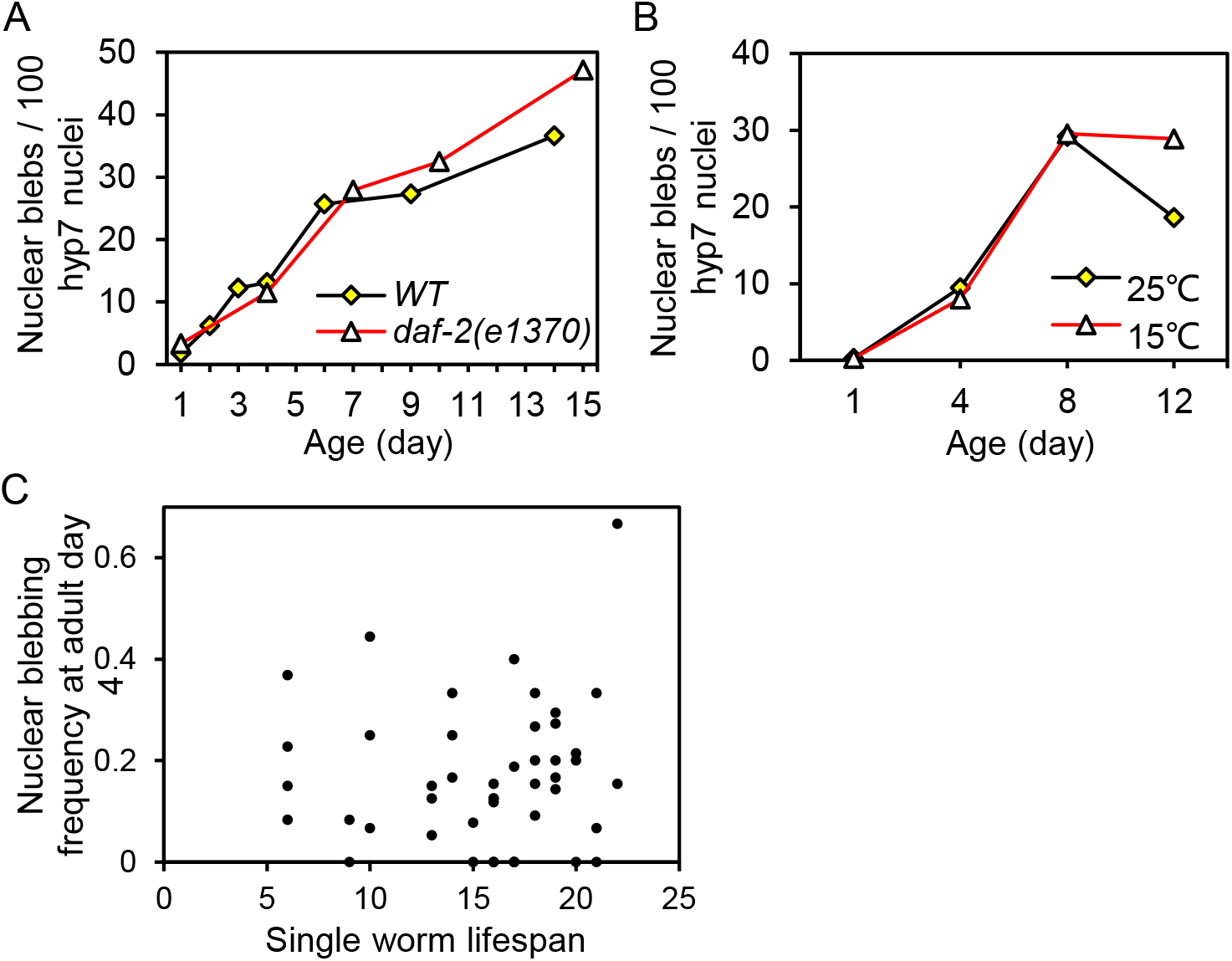
Means that alter the rate of aging have little or no effect on the frequency of nuclear blebbing. (A) The frequency of nuclear blebbing increased with age at the essentially the rate in the WT and the long-lived *daf-2(e1370)* mutant worms. Worms were maintained under the standard culture temperature of 20 °C. (B) WT worms cultured at 15 °C, which extends lifespan, and those cultured at 25 °C, which shortens lifespan, displayed the same nuclear blebbing frequency from AD 1 to AD 8. (C) No correlation between the lifespan of individual worms and the nuclear blebbing frequency measured on AD4. Spearman correlation, *r* = 0.115, *p* = 0.4468, two-tailed.

In addition, there are significant inter-individual differences in the rate of aging. Even for a population of isogeneic worms cultured in the same environment, e.g. on the same plate, the lifespans of individual worms vary dramatically (Kinser et al., 2021). We found that among the wild-type worms kept under the same condition, the frequency of nuclear blebbing at adult day 4 did not correlate with lifespan (Fig 4C).

Thus, the results above suggest that in C. elegans, the frequency of nuclear blebbing is not coupled with the rate of aging.

### 3.5 Proliferating germline stem cells promote nuclear blebbing in hypodermal cells

Since nuclear blebbing is uncoupled from aging (Fig 4) and is not induced by heat, starvation, and ROS (Fig 1C), we wondered what causes nuclear blebbing. We found that nuclear blebbing in the hypodermis responds to the activity of the gonad. The temperature-sensitive *glp-1(e2141)* mutation causes germ cells to enter meiosis prematurely at the restrictive temperature of 25 °C, resulting in an empty gonad of few germs cells and sterility; when cultured at 15°C, the *glp-1(e2141)* mutant has an intact gonad and reproduces normally (Austin and Kimble, 1987; Priess et al., 1987). We noticed that for the *glp-1(e2141)* mutant, the frequency of nuclear blebbing in the hypodermis was markedly lower at 25 °C than at 15 °C (Fig 5A), and subsequent quantification revealed a six-fold difference (~5 and ~30 blebs per 100 hyp7 nuclei on AD5, respectively, at 25 °C and 15 °C) (Fig 5D). As a control, wild-type worms showed no significant difference in the frequency of nuclear blebbing between 25°C and 15°C (Fig 5D). This result indicates that the mechanism that regulates nuclear blebbing of somatic cells responds to the activity of the reproductive system, not the culture temperature.

**Fig 5.**
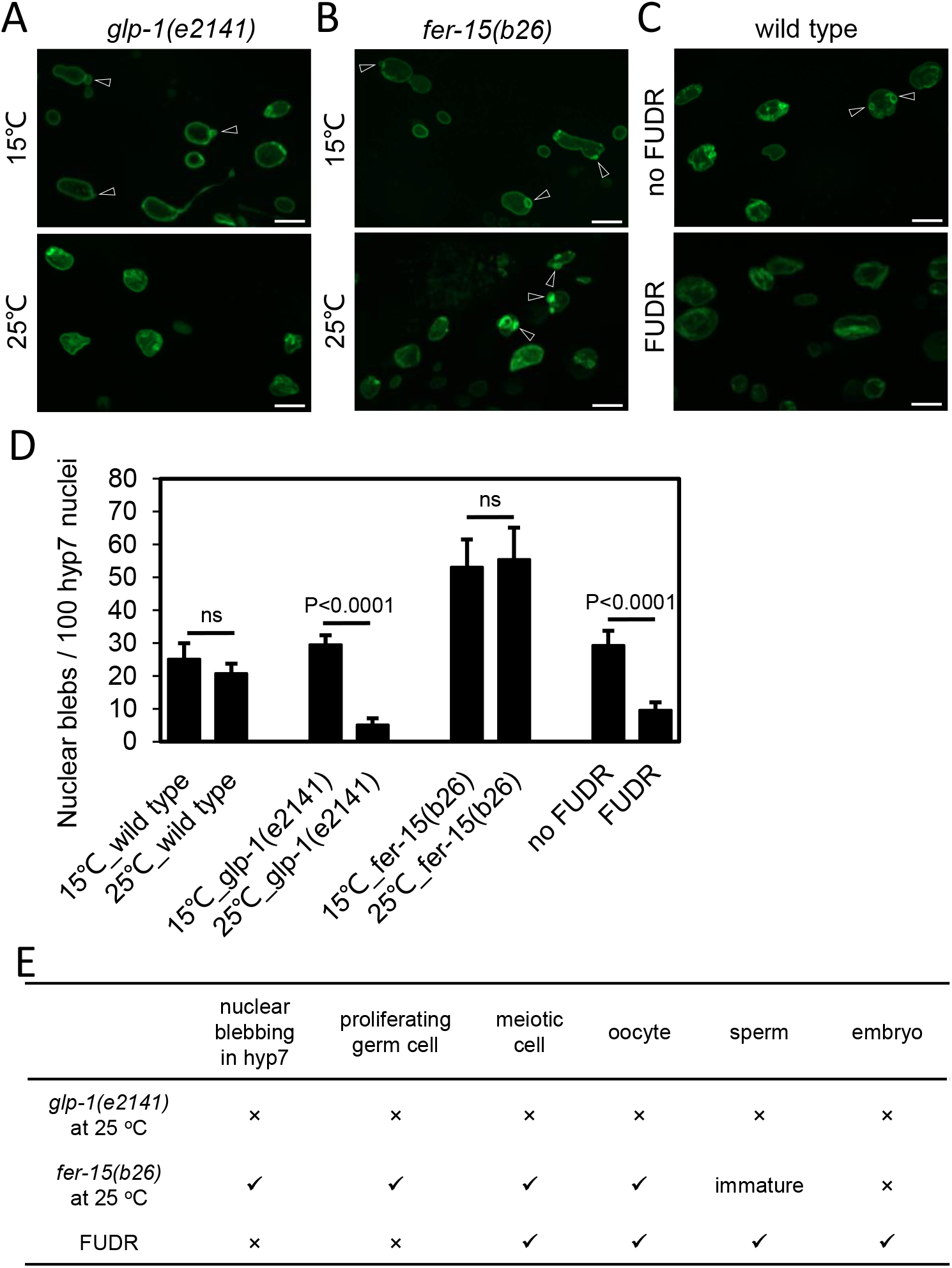
Proliferating germ cells promote nuclear blebbing in the hypodermis. (A, B, C) Representative images of nuclear blebs of *glp-1(e2141), fer-15(b26)*, and WT worms on AD5. (D) Nuclear blebbing frequency on AD5 under the indicated conditions. (E) Summary of the effects of FUDR, *glp-1(e2141)* at 25 °C, and *fer-15(b26)* at 25 °C on the germline and nuclear blebbing in the hypodermis. The *gfp::lmn-1* knock-in strain was used in (A-E). Scale bars, 10 μm. Arrowheads indicate nuclear blebs. Error bars represent standard error, and *p* values from t-test are shown. ns, not significant.

To further dissect the connection between somatic nuclear blebbing and the reproduction system, we analyzed the *fer-15(b26ts)* mutant, which is defective in spermatogenesis in a temperature sensitive manner (Hirsh and Vanderslice, 1976). The germ cells and oocytes of the *fer-15(b26ts)* worms are normal (Hirsh and Vanderslice, 1976). Intriguingly, there was no significant difference in the frequency of nuclear blebbing between *fer-15(b26ts)* worms cultured at the restrictive temperature of 25°C and those at the permissive temperature of 15°C (Fig 5B and 5D). Thus, nuclear blebbing in the hypodermis does not respond to the presence or absence of mature sperm, nor to embryos.

Next, we tested the effect of FUDR, a DNA synthesis inhibitor that can prevent the proliferation of germ cells (Golden et al., 2007). When FUDR treatment was initiated at mid L4 stage, we found that worms retained meiotic cells, oocytes, sperm and even embryos, but no proliferating germ cells (Fig S2 and 5E). Interestingly, the hyp7 nuclei of FUDR-treated worms accumulated fewer nuclear blebs than those of untreated worms (Fig 5C, 5D and S3).

Contrasting the above results, we find that the common denominator is evidently proliferating germ cells (Fig 5D). Proliferation of germ cells is inhibited in FUDR-treated WT worms and in *glp-1(e2141)* mutant worms cultured at 25 °C, and for both, hyp7 nuclear blebbing is also inhibited. Conversely, proliferation of germ cells is not affected in *fer-15(b26ts)* worms, neither is hyp7 nuclear blebbing (Fig 5C-D). Thus, we propose that proliferating germ cells promote nuclear blebbing in the hypodermis.

## 4. Discussion

### 4.1 Characteristics of nuclear blebbing in *C. elegans*

For more than half a century, the term “nuclear bleb” has been used to describe protrusions of the nuclear envelope into the cytoplasm under different biological conditions (Ahearn et al., 1967; Ahearn et al., 1974; Clark Jr, 1960; Dou et al., 2015; Furusawa et al., 2015; McDuffie, 1967; Mishra and Munnet, 1979; Ruddle, 1962; Szollosi and Szollosi, 1988; Törő and Olah, 1966; Vigouroux et al., 2001). However, these “blebs” differ in morphology, composition, cause of formation, and function. Our study showed that in aged *C. elegans*, nuclear blebs are enclosed by the nuclear lamina and nuclear envelope, and less than 50% of them contain chromatin. This is in line with literature reports. Depending on the biological context, nuclear blebs may be deficient in LamB (Furusawa et al., 2015; Vigouroux et al., 2001), NPCs (Karoutas et al., 2019; Vigouroux et al., 2001), or RNA Pol II (Karoutas et al., 2019), or enriched with γ-H2AX (Ivanov et al., 2013; Karoutas et al., 2019) or heterochromatin (Karoutas et al., 2019).

### 4.2 Nuclear blebbing contributes to chromatin loss through nucleus-to-cytoplasmic transport

*C. elegans* worms lose chromatin during aging (Fig S1 and (Golden et al., 2007)). Nuclear blebbing has been reported to mediate the nucleus-to-cytoplasm transport (Li et al., 2016; Patterson et al., 2011; Speese et al., 2012). Further considering that 42% of the nuclear blebs in aged *C. elegans* contain chromatin (Fig 2E), a fraction of which detach (Fig 3A), and detached blebs are targeted to lysosomes (Fig 3B-C), we reason that nuclear blebbing contributes to chromatin loss during aging. Other biological processes, such as nuclei loss, may also play a part (Golden et al., 2007; McGee et al., 2011).

With respect to the nucleus-to-cytoplasm transport via nuclear blebbing, one may ask what cargos the chromatin-negative blebs (Fig 2E) may contain. Our electron microscopy analysis did not reveal RNP-like granules (Li et al., 2016) in nuclear blebs of hypodermal cells. Given that Thioflavin T treated worms exhibited fewer blebs (Fig S4), whether some nuclear blebs are enriched in aggregated proteins is an interesting question.

### 4.3 Proliferative germ cells regulate nuclear blebbing in the hypodermis

Surprisingly, inhibiting proliferation of germ cells prevented nuclear blebbing in hypodermal cells (Fig 5). This observation echoes the previous findings that after worms reaching sexual maturity, germline stem cells abruptly turns down the heat shock response of somatic tissues (Labbadia and Morimoto, 2015; Shemesh et al., 2013). The regulatory role of the reproductive system is particularly fascinating when considering the disposable soma theory of aging, which suggests that aging is due to reallocation of resources to maximize reproduction at the expense of reduced error regulation in somatic cells (Kirkwood, 1977). Consistent with this idea, our observations support a model in which the reproductive system contributes to somatic aging through promoting nuclear blebbing and chromatin loss in somatic cells.

### 4.4 Nuclear blebbing and lamina convolution are regulated by distinct mechanisms

We propose that in aged *C. elegans*, nuclear blebbing and convolution of nuclear lamina are regulated by distinct mechanisms, based on the observations that: (1) convolution was slowed down in the *daf-2(e1370)* mutant, but blebbing was not; (2) inhibiting proliferation of germ cells prevented blebbing, but did not prevent convolution. Uncoupling of blebbing and other abnormal morphologies is not restricted to aged *C. elegans*. Previously, Chen et al. showed that fibroblasts lacking LamB1 or both LamB1 and LamB2 acquired more nuclear blebs but showed no changes in overall nuclear shape, whereas fibroblasts lacking all lamin genes (lamA/B1/B2) had more irregularly shaped nuclei but no change in nuclear blebbing frequency (Chen et al., 2018). Also, Coffinier et al. found that in cortical neurons of mice, LmnB1 deficiency induced nuclear blebbing, but LmnB2 deficiency led to elongation of nuclei instead of blebbing (Coffinier et al., 2011). Thus, different types of abnormalities in nuclear shape may reflect distinct biological processes.

## 5. Conclusions

In summary, we find that in *C. elegans*, the frequency of nuclear blebbing in hypodermal cells is not correlated with the rate of aging but is affected by proliferative germ line stem cells. Our findings suggest that somatic nuclear blebbing is not a biomarker of organismal aging in *C. elegans*.

## Authors’ contributions

Q.F., X.-M.L., S.-T.L. and M.-Q.D designed the experiments and interpreted the data. Q.F. performed most of the experiments. S.-T.L. performed the imaging experiments related to lysosome marker. C.Z. performed the EM analysis. Q.F. and M.-Q.D. drafted and revised the manuscript. M.-Q.D. supervised this study.

## Data availability

The data that support the findings of this study are available from the corresponding author upon reasonable request.

## Acknowledgements

We thank the Caenorhabditis Genetics Center (CGC), which is supported by the NIH Office of 628 Infrastructure Programs (P40 OD010440), for providing worm strains. We also thank Drs. Daniel A Starr, Xiaochen Wang and Xiao Liu for providing worm strain. This work was funded by National Natural Science Foundation of China (NSFC-ISF 32061143020 to M.-Q.D.), Ministry of Science and Technology of China (a fund of the National High-Level Talents Special Support Program to M.-Q.D. and institutional grants to NIBS, Beijing), and Beijing Municipal Science and Technology Commission (a fund for cultivation and development of innovation base to M.-Q.D. and institutional grants to NIBS, Beijing).

---

**Fig S1.**
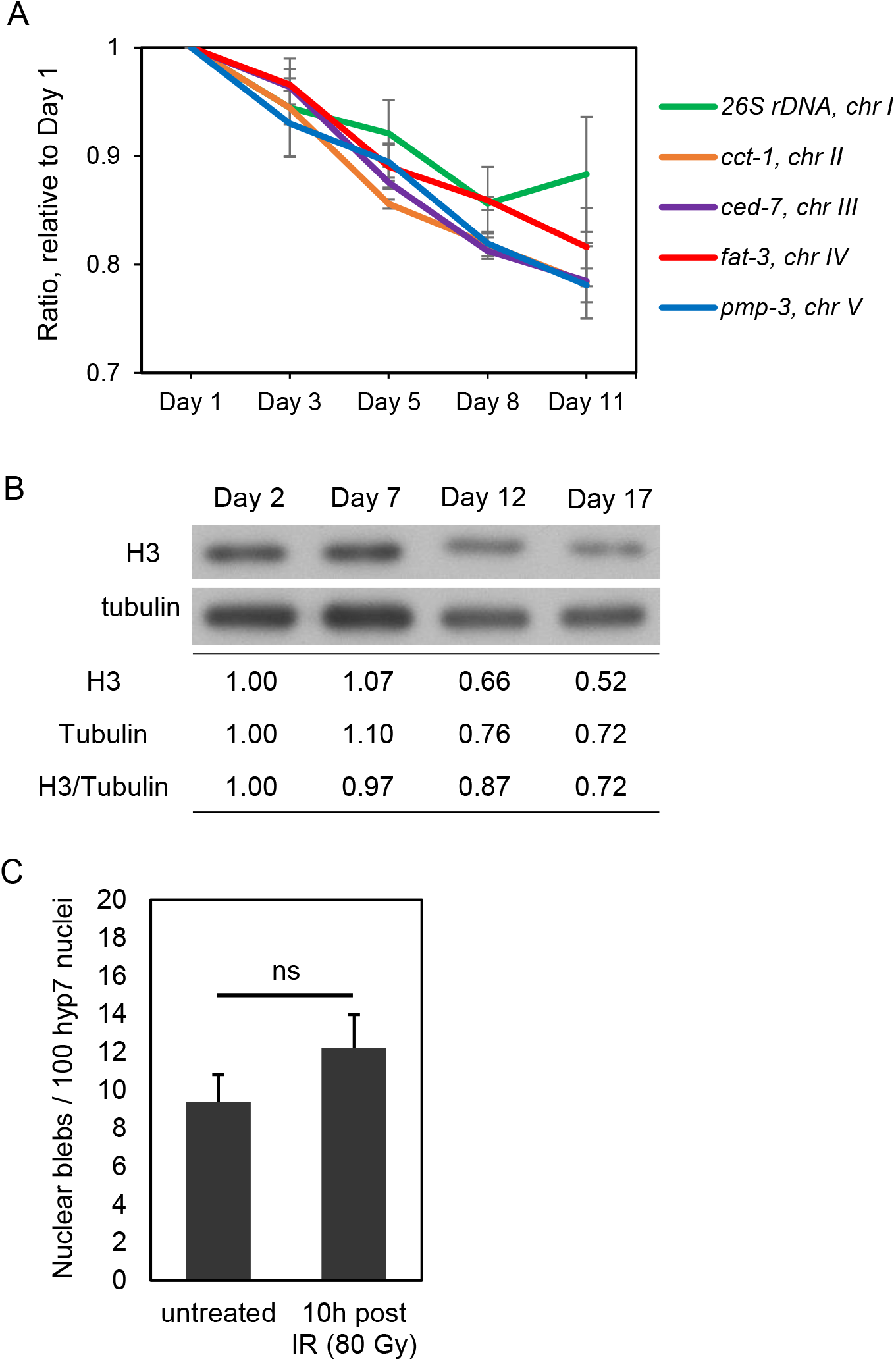
related to Fig 3. Chromatin loss during aging. (A) qPCR results verifying that gene copy numbers decreased during aging as reported previously (Golden et al., 2007). The data shown are the mean ± standard error from two independent biological replicates. (B) The level of histone H3 decreased during aging. The same number of worms were used for each sample. (C) γ-irradiation (IR) did not induce nuclear blebbing. ns, not significant, as determined by t-test.

**Fig S2.**
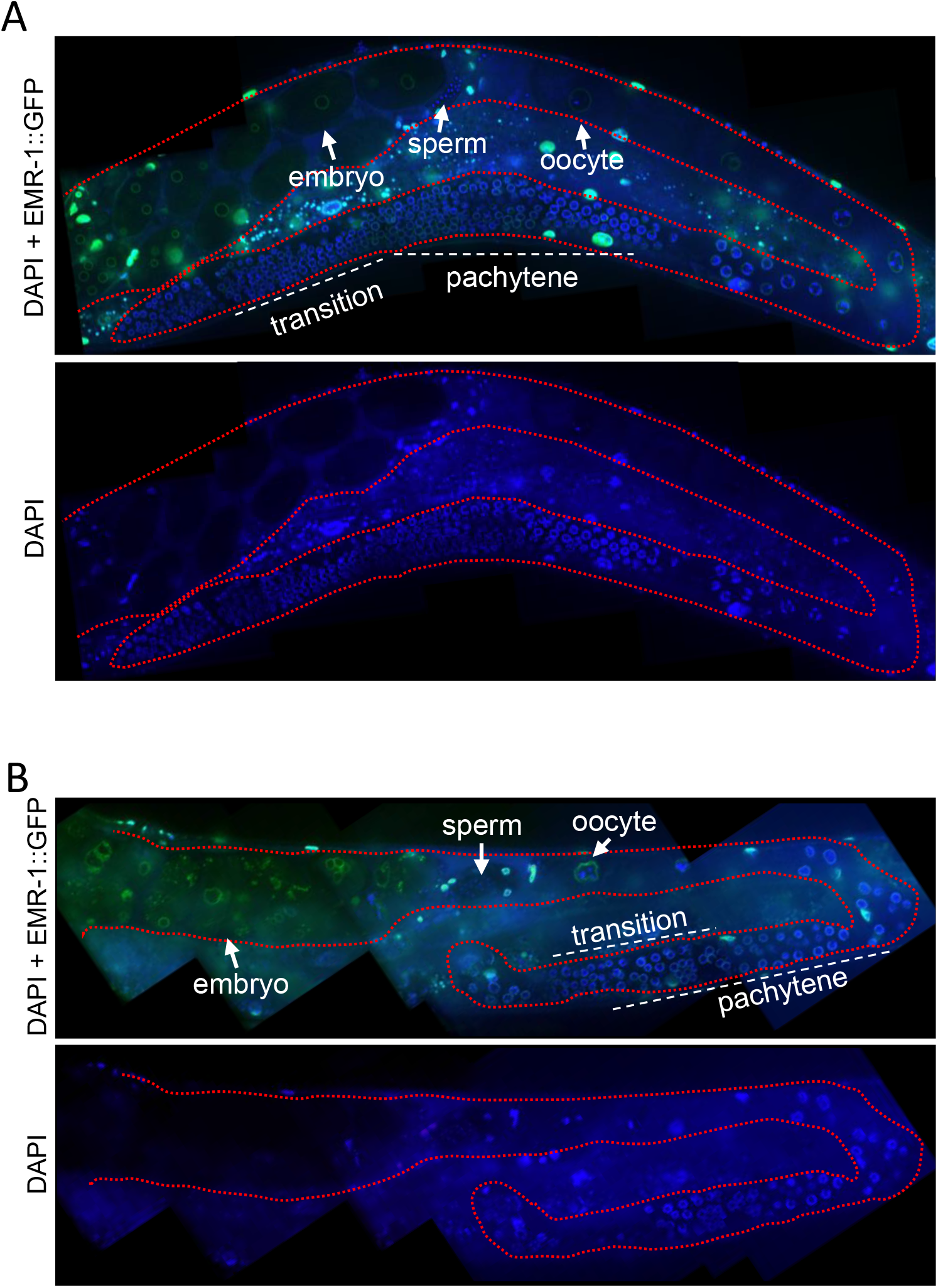
FUDR-treated worms have no proliferating germline stem cells. (A) Germline of an untreated control worm. (B) Germline of a FUDR-treated worm. The presence of meiotic cells, oocyte, sperm and embryo, and the absence of proliferating stem cells are shown. Scale bars, 10 μm.

**Fig S3.**
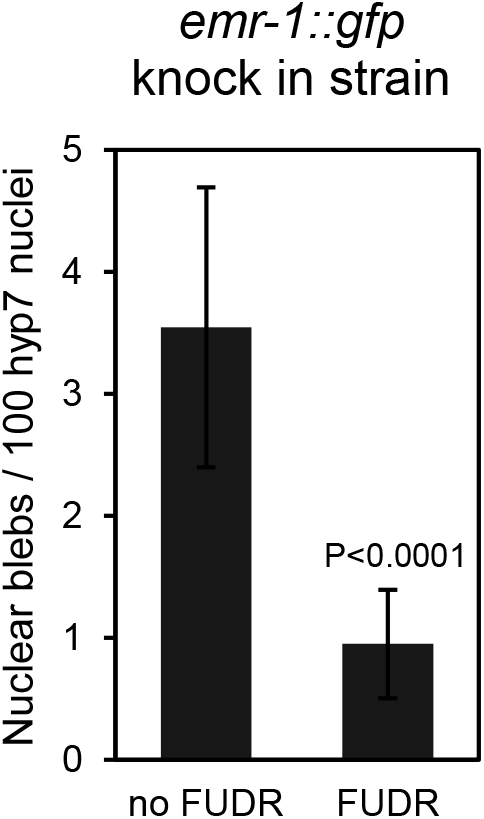
related to Fig 5. Nuclear blebbing is inhibited upon FUDR treatment in emr-1::gfp knock in strain. Nuclear blebbing frequency on AD5 in the *emr-1::gfp* knock-in strain.

**Fig S4.**
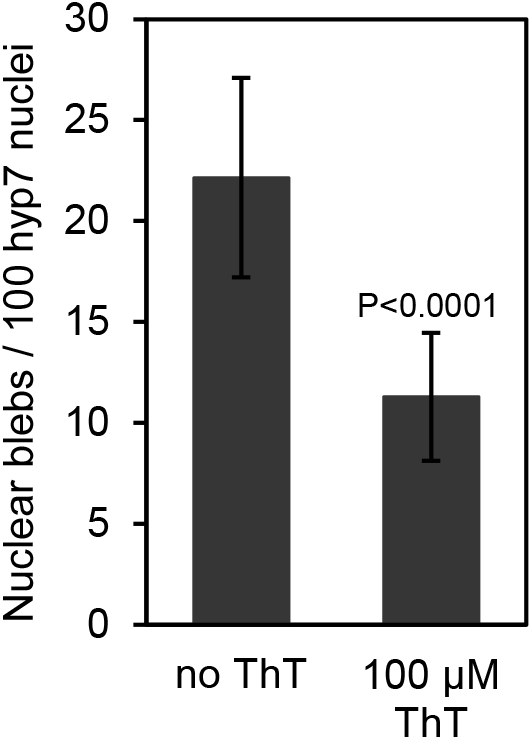
ThT-treatment ameliorates nuclear blebbing. On AD5, nuclear blebbing frequency was much lower in ThT treated worms. The hyp 7 nuclei of the *gfp::lmn-1* knock-in strain UD484 were imaged and analyzed

